# A Microgel-Based Platform for Tunable Expansion and Function of γδ T-cells

**DOI:** 10.64898/2026.07.23.740340

**Authors:** Favour Omafuvwe Obuseh, Junzhe Lou, Michelle Chang, Anqi Chen, David Weitz, David Mooney

**Affiliations:** Harvard-MIT Program in Health Sciences and Technology (HST), Cambridge, MA, USA; School of Engineering and Applied Sciences (SEAS), Harvard University, Cambridge, MA, USA; Wyss Institute for Biologically Inspired Engineering, Harvard University, Boston, MA, USA

**Author notes:** Corresponding Author. **Email:** David Mooney. These authors contributed equally to this work (Favour Omafuvwe Obuseh, Junzhe Lou). **Author Contributions:** Author contributions: Conceptualization: FO, JL, DM, Methodology: FO, JL, Investigation: FO, JL, MC, Visualization: FO, JL, Funding acquisition: FO, JL, DM, Writing – original draft: FO, JL, Writing – review & editing: FO, JL, MC, AC, DW, DM. **Competing Interest Statement:** Authors declare that they have no competing interests.

**Keywords:** γδ T-cells, Biomaterials, Immuno-engineering

## Abstract

Current γδ T-cell expansion protocols often sacrifice functionality for yield and largely ignore the context of activation. Here we utilize a tunable alginate microgel system functionalized with anti-CD3 and co-stimulatory antibodies (αCD28 or αCD2) to investigate the impact of biochemical signaling and substrate mechanics on γδ T-cell activation. Microgel-mediated expansion was compared to conventional soluble antibodies and TransAct beads. The microgels enhanced γδ T-cell expansion compared to soluble antibodies, allowed for controlled tuning of differentiation state, and promoted higher NKG2D, IFN-γ and TNF-α expression levels. Functionally, microgel-expanded γδ T-cells exhibited superior cytotoxicity against both solid and liquid tumor targets. This system also allowed elucidation of the differences in stimulation requirements for various donors, based on the starting phenotype. These findings establish a tunable platform for engineering γδ T-cells with improved therapeutic potential.

**Significance Statement:** γδ T-cells have shown promising therapeutic effects when used for T cell-based immunotherapy to treat solid tumor. However, achieving rapid expansion of γδ T-cells while maintaining their functionality remains a major challenge, especially given the heterogeneous responses from donors. We demonstrate that a tunable microgel system with flexible presentation of stimulatory cues improves γδ T-cell expansion while preserving cytotoxic function and reveal how starting phenotypes influence responses to activation. These understandings will provide design rationale to enable patient-specific treatment for optimal therapeutic outcomes.

## Introduction

T-cell based immunotherapy that leverages the specificity and toxicity of T-cells to recognize and clear tumor cells has proven to be an effective treatment for certain cancers(1–3). As a unique subset of T-cells, γδ T-cells have gained increasing attention in recent years and are now considered promising agents for the next generation of cellular immunotherapies(4–6). Unlike conventional αβ T-cells that are commonly used for T-cell immunotherapy, γδ T-cells, do not rely solely on classical MHC class I-mediated antigen presentation. Instead, they also utilize a range of innate-like receptors, including Natural Killer Group 2, member D (NKG2D) to recognize and eliminate stressed or malignant cells expressing ligands such as UL16-binding protein 1 (ULBP1), and MHC class I chain related protein A and B (MICA, and MICB)(6–8). These interactions trigger cytotoxic responses, including TNF-α mediated apoptosis, independent of MHC presentation(9, 10). This MHC-independence makes γδ T-cells particularly appealing for targeting tumors which shed MHC I, while also supporting the potential for allogenic use (6, 7, 10, 11). To further apply γδ T-cells for therapeutic treatments and potentially overcome challenges in cancer immunotherapy, efficient and rapid expansion of γδ T-cells with appropriate phenotypes and functions is required.

Recent advances have enabled the expansion of γδ T-cells, with strategies using soluble biologics, such as optimized cytokine cocktails (e.g., IL-2, IL-15, IL-18, IFN-γ) and soluble activating antibodies (αCD2, αCD3, αCD28), and in some cases feeder cells(5, 12–15). Soluble activating antibodies allow for facile activation protocols, with low TCR crosslinking preventing activation induced cell death. Although the use of soluble antibodies can result in over 100-fold expansion in 21 days(5, 13), the absence of strong TCR crosslinking, and or co-localization of co-stimulatory molecules does not capture the physiological processes relevant for γδ T-cell activation, where activating ligands are highly expressed on the surface of target cells in a coordinated manner(16–18).

Biomaterials have served as artificial cells that mimic native biological systems to activate and regulate immune cells, such as αβ T-cells(19–27). In these materials, surface ligand presentations and other biophysical features, such as stiffness and ligand mobility, can be tailored to modulate the phenotypes and functions of immune cells. However, γδ T-cell activation using biomaterial systems has been underexplored. Additionally, the specific role of stimulation cues in the differentiation of human peripheral blood γδ T-cells after thymic egress remains undetermined (7, 28), even though the higher basal γδ TCR density suggests a different, and perhaps more sensitive, response to cues (16, 29, 30), compared to αβ-T-cells (18, 29, 31, 32). Moreover, the composition of the starting T-cell population has been shown to significantly impact therapeutic outcomes. Patient-derived T-cells often underperform relative to healthy donors (33), and baseline phenotypic heterogeneity introduces further variability(34, 35). Accommodation for such differences requires personalized the T-cell activating dose. Altogether, these insights highlight the growing importance of designing highly flexible systems that allow modulation of various properties to account for the chemical and biophysical context of immune activation, in order to produce functional γδ T-cells.

Here we developed an alginate-based microgel system with tunable properties to control γδ T-cell activation, expansion, and cytotoxic potential, and to investigate the role of various biological cues. Upon precise modulation of various material features, we demonstrate that expansion, activation phenotypes and cytotoxic functions are regulated by activating antibody density. We also show greater expansion with microgels, as compared to soluble antibodies, and increased cytotoxicity of expanded cells when compared to the commercially available TransAct system. Furthermore, tuning microgel parameters can optimize γδ T-cell products, while resolving the starting donor variability that exists even across healthy donors.

## Results

### Fabrication of Microgel System for Expansion of γδ T-Cells

Alginate microgels with covalent crosslinks were fabricated via microfluidic emulsion by mixing norbornene and tetrazine modified alginate at a polymer concentration of 2 wt%, yielding microgels with a diameter of 77 ± 2 μm (Fig 1a, (19). The elastic modulus of microgels can be modulated by varying the degree of substitution of norbornene and tetrazine groups on alginate polymers (Fig.S1). To engineer these microgels as artificial antigen-presenting cells for γδ T-cell expansion, microgels were then surface functionalized with essential stimulatory antibodies using a previously described layer-by-layer approach(19, 36). Briefly, alginate microgels were first soaked in a solution of poly(D-lysine) to surface coat with a layer of positively charged polymers and then immersed in solution of azide-labelled alginate to deposit another layer of negatively charged polymers for further modification (Fig. 1a-c). Various stimulatory antibodies modified with dibenzocyclooctyne (DBCO) were then efficiently conjugated to microgel surface via azide-DBCO click chemistry. This method provides a convenient means to precisely engineer the surface biochemical properties independent of microgel stiffness. These polymers coating layers are stable in culture media over weeks, as demonstrated in our previous work (19).

**Figure 1.**
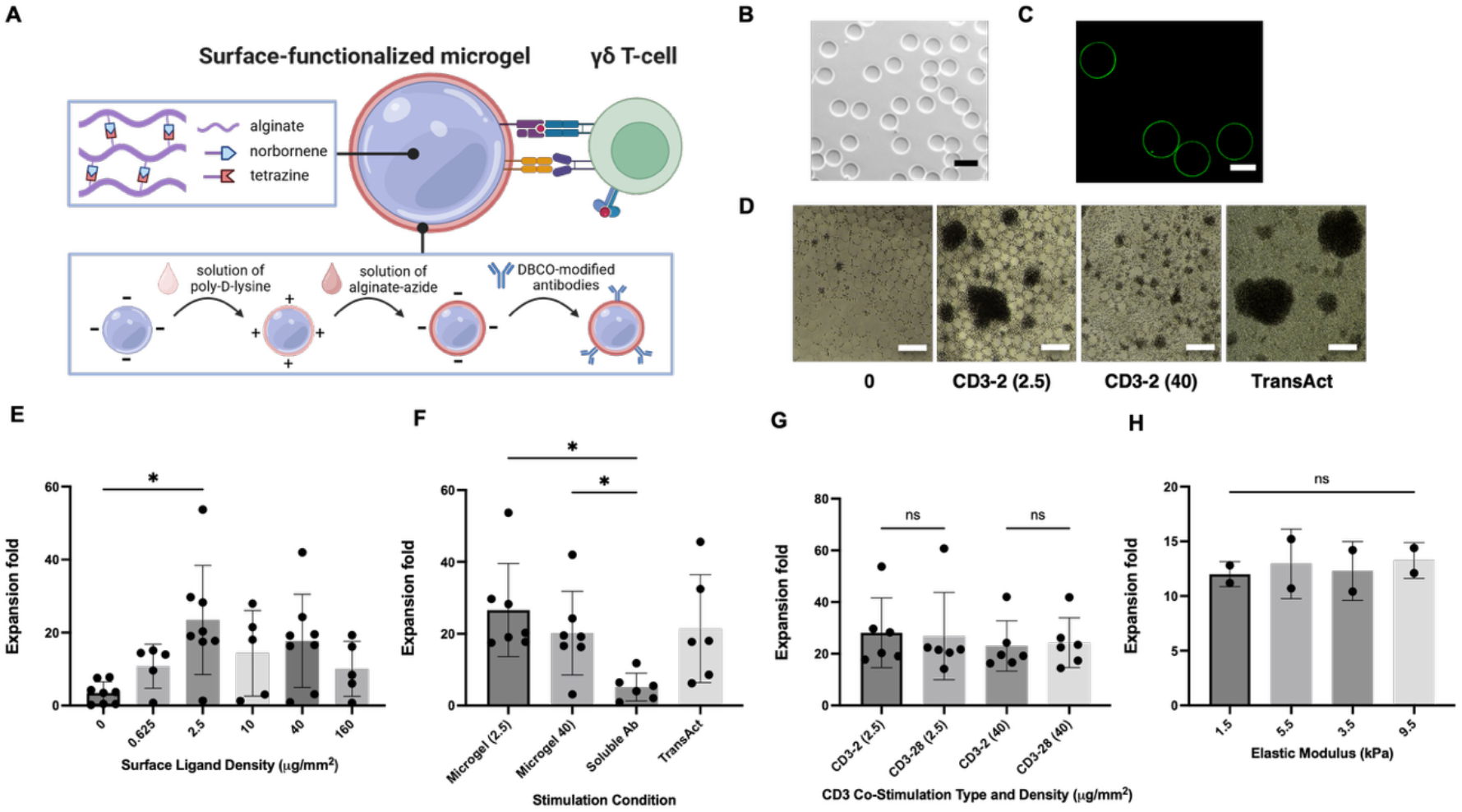
Fabrication of a microgel system with tunable properties for γδ T-cell expansion. (A) Schematic of surface-functionalized alginate microgels presenting stimulatory antibodies. Alginate-based microgels are crosslinked via norbornene–tetrazine click chemistry, and microgel surfaces are modified using a layer-by-layer coating strategy with poly(D-lysine) and alginate to enable antibody presentation. (B) Representative phase-contrast image of alginate microgels. Scale bar: 100 μm. (C) Confocal images of fluorescently labelled alginate coated on the surface of microgels. Scale bar: 50 μm. (D) Representative phase-contrast images of primary human γδ T-cells cultured with blank microgels, αCD3/αCD2-functionalized microgels, or TransAct. (E) Expansion of primary human γδ T-cells as a function of microgel surface antibody density on day 9. (F) Expansion of primary human γδ T-cells using antibody-functionalized microgels compared with soluble antibodies and TransAct. Scale bar, 200 μm. (G) Expansion of primary human γδ T-cells as a function of costimulatory ligand (CD2 versus CD28) on day 9. (H) Expansion of primary human γδ T-cells as a function of microgel stiffness on day 9. For panel E,F,G, n = 5–8 donors. H, n = 2 donors. Bar graphs show mean ± s.d. Statistical significance was determined using ordinary one-way ANOVA with Tukey’s multiple comparisons test where applicable. Prism mixed-effects model analysis was used for datasets with missing values (where ordinary two-way ANOVA could not be performed). *P < 0.05, **P < 0.01, ***P < 0.001, ****P < 0.0001; ns, not significant.

To evaluate the efficacy of the microgel system for activating and expanding γδ T-cells, we first systematically varied overall surface antibody density from 0-160 ug/mm^2^. In these studies, a 2:1 ratio of αCD3: αCD2 antibodies was employed on the microgel surface, as inspired by previous work using soluble antibodies to expand γδ T-cells with high cytotoxic potential(13). The microgels were co-cultured with human γδ T-cells in the presence of IL-15 and IL-18, cytokines known to enhance cytotoxicity potential of γδ T-cells. We observed that increasing surface antibody density initially led to greater fold expansion of γδ cells; however, expansion decreased above a threshold of 2.5 μg/mm^2^. In parallel, we observed a decrease in cell cluster size above the threshold. (Fig 1d,e). Compared to conditions using soluble antibodies for activation, the microgel platform induced superior expansion, reaching levels comparable to commercially available TransAct nano-polymeric beads (Fig 1f).

We next explored the role of co-stimulatory antibodies by comparing cell expansion using αCD2 and αCD28 as stimulatory ligands (Fig 1g). No significant difference was observed in expansion, across two different overall density conditions (40 ug/mm^2^ and 2.5 ug/mm^2^). We further investigated the impact of microgel stiffness on γδ T-cell expansion. Microgels with stiffness ranging from 1.5 -9.5 kPa were functionalized with αCD3/αCD2 antibodies at 2.5 ug/mm^2^ and co-cultured with γδ T-cells. In contrast to reports with αβ T-cells that are sensitive to the mechanics of stimulatory materials(19, 37), the fold expansions of γδ T-cells were similar when using microgels of different stiffness (Fig 1h).

As these studies revealed that ligand density plays a critical role in activation and expansion of γδ T cells, subsequent experiments focused on investigating how ligand density affects γδ T-cell phenotype and function. αCD3**/**αCD2 antibodies at an overall stimulation density of 2.5 ug/mm^2^ (low-density) and 40 ug/mm^2^ (high-density) were chosen as activation conditions, due to their ability to induce substantial T cell expansion.

### Surface Antibody Density Modulates γδ T-Cell Phenotype and Activation

To initially investigate how ligand density affects differentiation states of activated γδ T-cells, expanded cells were characterized for CD27 and CD45RA surface expression. These have been widely used to delineate γδ T-cell subsets based on their proliferative capacity and association with terminal differentiation(34, 35, 38). While it remains debated which subset is most beneficial in vivo(13, 35, 39), CD27^−^ CD45RA^**+**^ cells are generally associated with cytotoxic effector function and terminal differentiation(13, 35, 39). Microgel-mediated expansion led to an increased proportion of CD27^−^ CD45RA^+^ γδ T-cells at the lower stimulatory antibody density (2.5 ug/mm^2^), alongside preservation of the CD27^+^ CD45RA^+^ population within the Vδ1 subset (Fig 2a). The latter is associated with a less differentiated/naïve-like phenotype. In contrast, the higher surface antibody density (40 ug/mm^2^) led to an increase in the CD27^-^ CD45RA^-^ population, which is typically associated with effector memory T-cells (Fig 2a,b). In Vδ2 cells, the lower antibody density similarly led to a higher frequency of CD27^−^ CD45RA^+^ cells, whereas higher densities favored differentiation toward the CD27^−^ CD45RA^**−**^ phenotype, mirroring patterns observed with TransAct expansion (Fig 2a,c).

**Figure 2.**
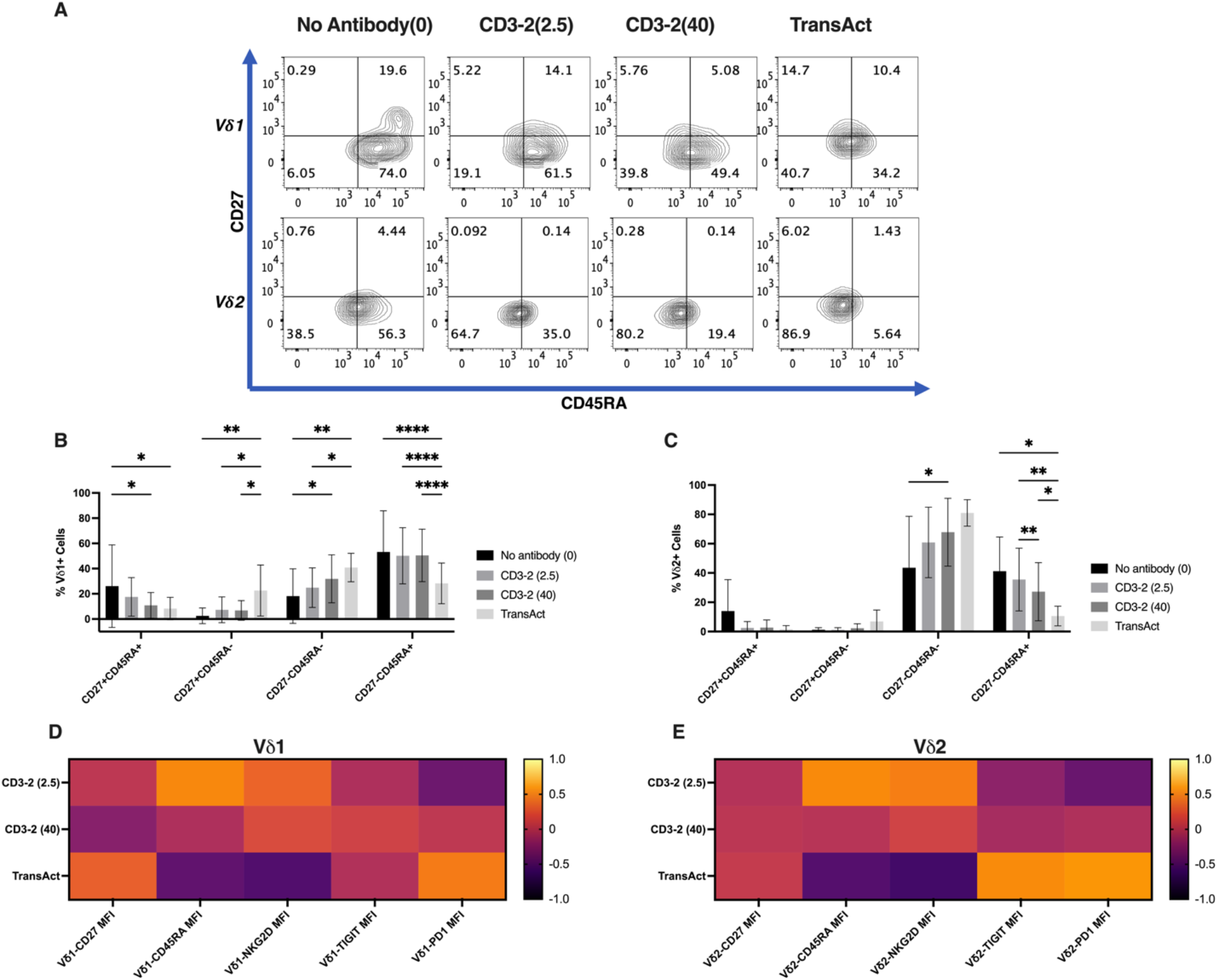
Surface Antibody Density Modulates γδ T-Cell Phenotype and Activation. (A) Representative flow cytometry contour plots of CD27 vs. CD45RA expression in expanded Vδ1 (top row) and Vδ2 (bottom row) for γδ T-cells cultured on microgels with varying surface antibody densities, using a αCD3/αCD2 microgel platform compared to unmodified gel without surface antibodies (Blank), and commercially available TransAct. (B,C) Quantification of differentiation profiles within the Vδ1, Vδ2 compartment respectively, (D,E) Heatmap of z-score– normalized geometric mean fluorescence intensity (gMFI; scale shown) for Vδ1 and Vδ2 cells respectively, across conditions for CD27, CD45RA, NKG2D, TIGIT, and PD-1. For panels B–C, n = 8 donors, D,E, n=9, 10 donors bar graphs represent mean ± s.d. Prism mixed-effects model analysis was used for datasets with missing values (where ordinary two-way ANOVA could not be performed). *P < 0.05, **P < 0.01, ***P < 0.001, ****P < 0.0001; ns, not significant.

Consistent with the differentiation shifts for the Vδ1 compartment, NKG2D geometric mean fluorescence intensity of the total population (gMFI) increased inversely with stimulation density, and remained higher across all microgel conditions as compared to TransAct-stimulated cells (Fig. 2d). PD-1 gMFI showed a modest increase with higher surface antibody density and was elevated in TransAct-stimulated cells. While TIGIT gMFI did not differ significantly across antibody densities overall (Fig. 2d), a density-dependent difference in the percentage of TIGIT^+^ cells was observed when comparing low versus high surface antibody stimulation (Fig.S2a,b).

For the Vδ2 cell population, NKG2D gMFI similarly increased at lower stimulation densities, and was higher across microgel conditions as compared to TransAct-stimulated cells (Fig. 2e). In similar fashion to Vδ1 cells, TIGIT percentages differed significantly between low and high-surface-antibody density stimulation, and PD-1 gMFI again showed only a modest increase with increasing surface antibody density but was elevated in TransAct-stimulated cells (Fig. 2e; Fig. S2c,d). Together, these data suggest that modulating signal strength through surface antibody density can balance cytotoxic-associated activation with less differentiated phenotypes during ex-vivo expansion, without leading to the higher inhibitory receptor levels observed under TransAct expansion.

### Microgel-Based Activation Enhances γδ T-Cell TNF-α/IFN-γ expression and Cytotoxicity

To assess the functional capacity of γδ T-cells expanded using the microgel platform, intracellular cytokine staining was performed to quantify effector responses. γδ T-cells expanded using either the microgel platform or TransAct were stimulated with phorbol myristate acetate (PMA) and ionomycin to induce maximal cytokine production, and intracellular effector markers were subsequently measured by flow cytometry. Compared to the TransAct-expanded cells, the percentage of TNF-α^+^IFN-γ^+^ cells was much higher in microgel-expanded cells (Fig. 3a), indicating a more cytotoxic phenotype. This was consistent with elevated expression of IFN-γ and TNF-α for Vδ1 and Vδ2 cells, respectively, as compared to TransAct expansion (Fig. 3b–e). The percentage of Granzyme B^+^IFN-γ^+^ cells was also elevated when compared to TransAct expansion (Fig. S3).

**Figure 3.**
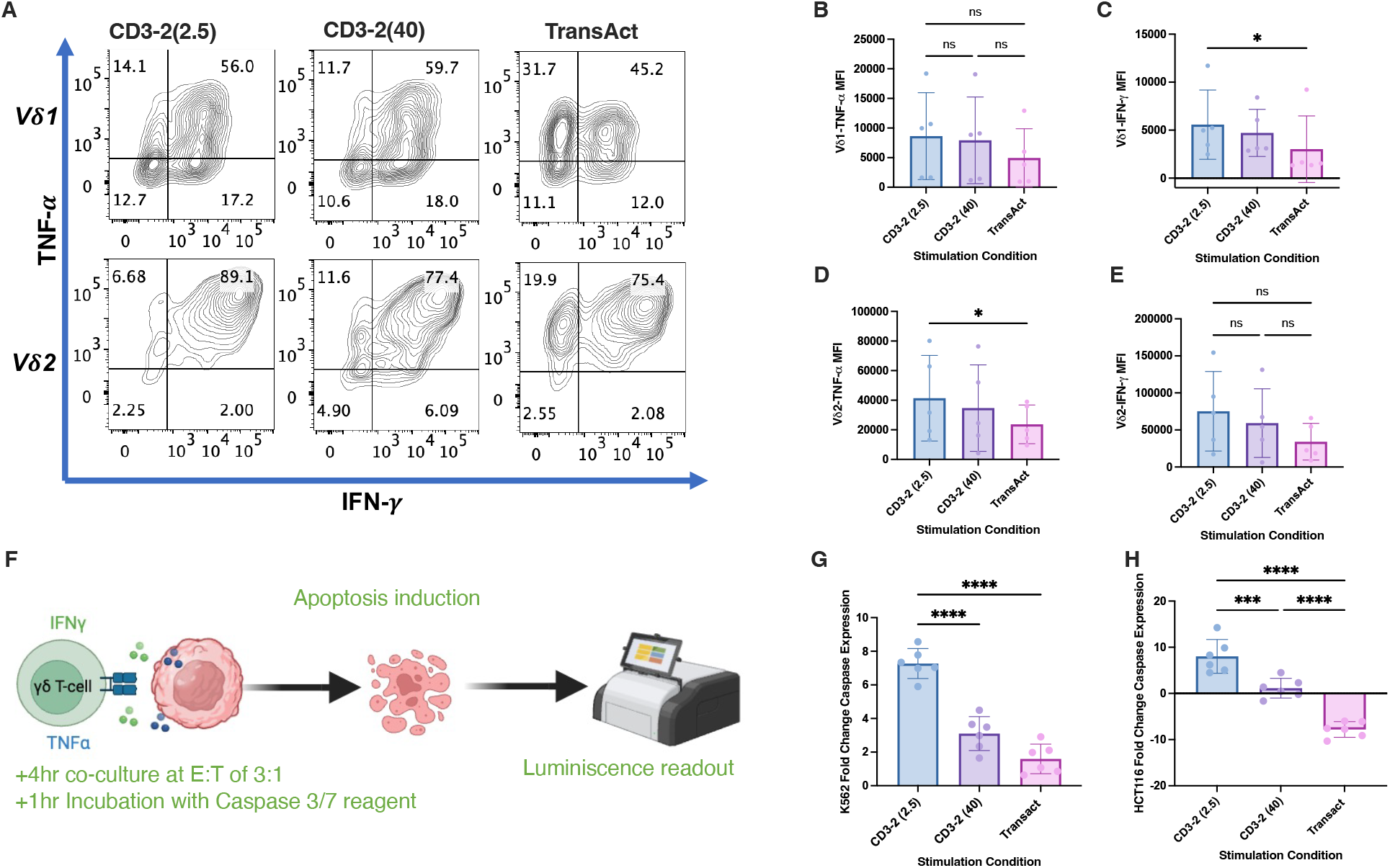
Microgel-Based Activation Enhances γδ T-Cell TNF-α/IFN-γ expression and Cytotoxicity. (A) Representative flow cytometry contour plots of TNF-α versus IFN-γ in expanded Vδ1 (top row) and Vδ2 (bottom row) γδ T-cells following culture on αCD3/αCD2-functionalized microgels at varying surface antibody densities, compared with TransAct. (B–C) TNF-α and IFN-γ geometric mean fluorescence intensity (gMFI), respectively, within the Vδ1 compartment across expansion conditions. (D–E) TNF-α and IFN-γ gMFI, respectively, within the Vδ2 compartment across expansion conditions. (F) Schematic of the caspase-3/7 apoptosis induction assay (created with BioRender). (G–H) Fold change in caspase-3/7 signal in K562 (G) and HCT116 (H) target cells following 4 h co-culture with γδ T-cells expanded under the indicated conditions. For panels B–E, n = 5 donors; for panels G–H, n = 6 technical replicates. Bar graphs show mean ± s.d. Statistics were performed using ordinary one-way ANOVA with Tukey’s multiple comparisons test. *P < 0.05, **P < 0.01, ***P < 0.001, ****P < 0.0001; ns, not significant.

We next examined the ability of expanded γδ T-cell to induce apoptosis of K562 leukemia cells when in co-culture (Fig. 3f). γδ T-cells expanded under the low surface antibody density condition exhibited markedly enhanced cytotoxic activity, compared to both the high surface antibody density expansion and TransAct-expanded cells (Fig. 3g). A similar trend was observed against the aggressive solid tumor cell line HCT116 (Fig. 3h), demonstrating a similar impact against both solid and hematologic tumor targets. The role of co-stimulation signals was also examined by substituting αCD28 for αCD2 in the microgels used for expansion. While the incorporation of αCD28 ligand did not significantly reduce expression of cytotoxic markers, it underperformed in tumor killing assays (Fig. S4a-h).

### Microgel-Based Activation Reveals Donor-Dependent Tuning

Throughout the previous analysis, substantial donor-to-donor variation was observed in γδ T-cell responses to stimulatory ligands. Donor-to-donor variability is a key challenge in T-cell immunotherapy, contributing to heterogeneity in manufactured products and variability in clinical response, including treatment failure in a subset of patients(40, 41). Leveraging our highly tunable material system, we sought to understand how initial donor cell phenotype impacted expansion outcome, and its dependency on activation parameters.

A review of the starting cells revealed three main donor starting differentiation phenotypes (Fig. 4a), which were manually grouped into γδ T-low, γδ T-intermediate, and γδ T-high based on the expression of markers associated with activation history (Fig. 4b). The γδ T-low group had lower average Vδ1/Vδ2 ratios and lower NKG2D, PD-1, and TIGIT percentages. The γδ T-intermediate group was characterized by higher average Vδ1/Vδ2 ratios and elevated NKG2D, PD-1, and TIGIT percentages. The γδ T-high group was characterized by the highest average Vδ1/Vδ2 ratios (∼8X that of the γδ T-low group), greatly diminished NKG2D percentage, and markedly increased PD-1 and TIGIT percentages. Overall, γδ T-high donors showed the highest expression of inhibitory-receptors.

**Figure 4.**
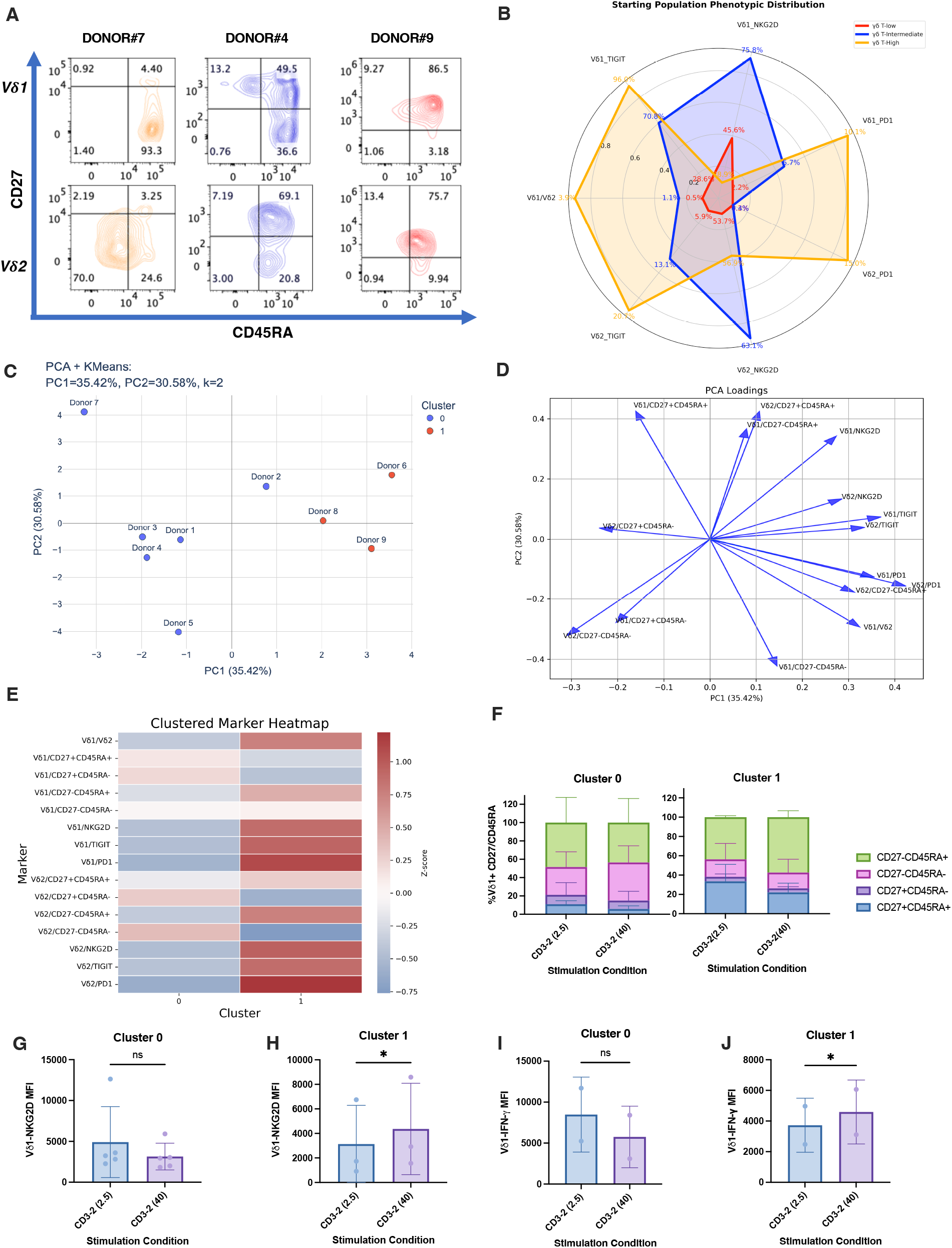
Microgel-based activation reveals donor-dependent tuning of activation-associated phenotypes. (A) Representative day 0 flow cytometry plots for each donor, highlighting baseline CD27 and CD45RA distributions within the Vδ1 and Vδ2 compartments. (B) Radar (spider) plot summarizing TIGIT, PD-1, NKG2D, and the Vδ1/Vδ2 ratio across donor composite-phenotype groups; both absolute values and normalized values (0.1–0.9) are shown. Donors were assigned to composite phenotype groups defined by inhibitory receptor expression (TIGIT, PD-1), cytotoxic receptor status (NKG2D), and subset composition (Vδ1/Vδ2 ratio): γδ T-low (n = 4; lowest TIGIT and PD-1), γδ T-intermediate (n = 4; high NKG2D and high TIGIT with intermediate PD-1), and γδ T-high (n = 1; highest PD-1 and TIGIT with the highest Vδ1/Vδ2 ratio). (C–E) Unsupervised post-expansion analysis. Principal component analysis (PCA) with k-means clustering of donor responses (n = 9) was performed using the difference between CD3-2 (2.5) and CD3-2 (40), after subtracting each donor’s day 0 baseline phenotype. The number of clusters was selected based on the highest average silhouette score. (D) PCA loading plot showing the relative contribution of each measured feature to PC1 and PC2. (E) Heatmap of scaled marker values with cluster assignments.(F) CD27/CD45RA quadrant frequencies within Vδ1 γδ T-cells for donors stratified by the unsupervised clusters (cluster 0 vs cluster 1).(G–H) NKG2D geometric mean fluorescence intensity (gMFI) within the Vδ1 compartment comparing low-density versus high-density expansion conditions for the two clusters.(I–J) IFN-γ gMFI within the Vδ1 compartment comparing low-density (CD3-2 (2.5) versus high-density (CD3-2 (40) expansion conditions for the two clusters. For panels G–J, sample sizes are indicated per panel (G, n = 5; H, n = 3; I, n = 2; J, n = 2). Bar graphs show mean ± s.d. Statistical comparisons between low- and high-density conditions were performed using Prism’s ratio paired t test (paired by donor). *P < 0.05, **P < 0.01, ***P < 0.001, ****P < 0.0001; ns, not significant.

We next analyzed how the post-expansion phenotype was impacted by the varying starting populations. The initial (pre-expansion) donor phenotype (baseline) was subtracted from the high (40 ug/mm^2^) vs low (2.5 ug/mm^2^) surface antibody expansion conditions, and the resulting delta was used for unsupervised clustering and k-means analysis to characterize phenotypic outcomes post stimulation. The clustering seed of 2 was determined based on the highest silhouette score. The donors classified as γδ T-low formed cluster 1, while the γδ T-high and γδ T-intermediate donors clustered together in 0 (Fig. 4c). The PC loadings showed that the driving force for cluster 0 was overall change in TIGIT percentages, coupled with NKG2D, PD-1, and the Vδ1/Vδ2 ratios (Fig. 4d). A cluster heatmap was also generated, highlighting the differences between both clusters, and demonstrating that cluster 1 donors were sensitive to increased stimulation dose, with overall activation increased at the higher dose (Fig. 4e).The higher dose stimulation resulted in a higher CD27^−^CD45RA^−^ fraction for cluster 0 donors, while leading to a higher CD27^−^CD45RA^+^ fraction for cluster 1 donors (Fig. 4f). With regards to activation and priming for cytotoxicity, an increase in NKG2D and IFN-γ expression was observed with higher stimulation for cluster 1 donors as compared to cluster 0 donors, which had the opposite trend (Fig. 4g–j).

## Discussion

Here we describe the use of a biomaterial system with tunable properties for the activation of γδ-T-cells. Leveraging this biomaterial system, we showed that surface antibody conjugation plays a critical role in expansion, while varying co-stimulation and materials stiffness had comparatively minor effects. The tunable surface density enables modulation of γδ-T-cell activation, cytotoxic marker expression, and target cell cytotoxicity. Importantly, the response to surface antibody density was found to be dependent on the donor starting population, with a higher surface antibody density beneficial for naïve-like donors, and a lower surface antibody density beneficial for more experienced donors.

Surface conjugation of αCD3/αCD2 to the microgels resulted in significant expansion of γδ-T-cells, when compared to soluble antibody-based expansion, but modulation of stiffness and co-stimulatory ligands had minimal effects. Covalently linking the antibodies to the surface of the microgels resulted in over a 4x difference in expansion when compared to soluble antibodies, with an average of 26-fold expansion observed over in just 9 days. Previously, the use of αCD3**/**αCD2 soluble antibodies was reported to yield ∼117-fold expansion over the course of 3 weeks(13). Unlike traditional expansion modalities, surface antibody conjugation provides controlled biochemical modulation, allowing for sustained, repeated receptor engagement as well as improved co-localization of both stimulatory and costimulatory ligands, which is likely beneficial for activating γδ-T-cells. When the co-stimulatory antibody was changed from αCD2 to αCD28, there were no significant observed changes in expansion and differentiation state. However, expansion with the αCD28 ligand resulted in decreased cytotoxicity of the resultant cell population. This suggests that the NF-κB–dominant signaling profile initiated by the αCD28 co-stimulation may be suboptimal for γδ T-cell tumor immunotherapy, in contrast with αCD2 co-stimulation, which preferentially engages NFAT-mediated signaling, supporting a more effective cytotoxic phenotype(42, 43). We further analyzed the impact of a range of microgel stiffness from 1.5-9.5 kPa on γδ T-cell activation. No significant impact was found with this range, in regard to expansion, and phenotype modulation. This is consistent with reports that γδ T-cell receptors (TCRs) may lack the force-dependent mechanosensing behavior described for αβ TCRs, where bioforces modulate bond lifetimes with peptide–MHC (pMHC) complexes(44). In contrast, γδ TCRs recognize a diverse array of antigens through mechanisms thought to be less dependent on mechanical engagement(44, 45). Nonetheless, additional work is warranted to evaluate mechanical effects in the context of encapsulation, as γδ T-cells are highly migratory and encounter a wide range of tissue microenvironments with distinct mechanical properties. Although tuning mechanical properties of the microgel system did not augment expansion and functionality, our system allowed for leveraging biochemical cues to modulate expansion of γδ-T-cells.

This microgel system with tunable surface density allowed for control over activation phenotype and cytotoxicity potential of the expanded cells. Changes in CD27/CD45RA population distribution, percentages and expression of NKG2D, TNF-α, and IFN-γ as well as, target cell cytotoxicity were observed as a function of antibody surface density. While a higher surface antibody density might be expected to drive terminal differentiation through stronger TCR signaling, our findings suggest that it instead promotes the accumulation of markers associated with an effector-memory–like phenotype (CD27^−^ CD45RA^−^ γδ T-cells). In contrast, lower surface antibody density microgels support a broader phenotypic landscape, permitting both the emergence of highly cytotoxic CD27^−^ CD45RA^+^ cells and the retention of marker expression associated with naïve like subsets (CD27^+^ CD45RA^+^), particularly in the Vδ1 compartment. One might expect that increasing the surface antibody density, which could mimic in-vivo high antigen load, would drive a more CD27^-^CD45RA^+^ responder population which is highly cytotoxic, at the expense of long-lived memory cells. However, the unexpected effects of surface antibody density highlight a possible difference in our understanding of γδ-T-cells vs αβ-T-cells and how they respond to increased stimulation density(19, 31). When evaluating functional response through cytotoxicity analysis, the cells from the lower density expansion with preserved naïve-like (CD27^+^ CD45RA^+^) and cytotoxic (CD27^−^ CD45RA^+^) phenotype showed higher expression of NKG2D, TNF-α, and IFN-γ, while achieving greater target-cell apoptosis against aggressive HCT116 cells, and K562 cells. The ability to tune the phenotype and cytotoxic potential of the cells suggest that the microgel system can be modulated to enrich beneficial populations in the context of cytotoxicity.

Although γδ T-cells expanded using microgels with low surface antibody density led to overall increased expansion and cytotoxic potential, further review of the response to stimulation showed strong dependence on donor starting populations. Donors with higher baseline Vδ1/Vδ2 ratios tended to exhibit increased frequencies of TIGIT^+^ and PD-1^+^ γδ T cells and a greater proportion of CD27^−^CD45RA^+^ cells, features consistent with more antigen-experienced or effector-skewed states. Donors enriched for CD27^+^CD45RA^+^ γδ T cells (less differentiated/”naive-like”) required higher αCD3**/**αCD2 signaling to transition into effector states and to mediate cytotoxicity, whereas antigen-experienced donors, enriched for CD27^−^CD45RA^+^effector-like populations, often with higher TIGIT expression and a higher Vδ1/Vδ2 ratio, achieved comparable responses at lower surface antibody density. The concordance across between starting populations provides supporting evidence that the phenotypic differences observed across donors reflect variation in prior antigen exposure rather than solely culture-induced effects. Notably, in some donors the Vδ2 compartment appeared more “primed” than the Vδ1 compartment, which likely contributed to subset-specific differences in stimulation sensitivity and functional output. In adults, it has been shown that enrichment of Vδ1 relative to Vδ2 is associated with antigen-driven reshaping of the γδ repertoire, consistent with the concept that immune history can shift baseline subset composition (34, 46). Our data suggests that donors which have phenotypes consistent with prior possible chronic activation, as evidenced through increased PD-1, TIGIT, and decreased NKG2D starting percentages, lower stimulation may be beneficial, as compared to more naïve-like donors which benefit from higher stimulation. These findings underscore the importance of considering both Vδ1 and Vδ2 starting phenotypes and their potential crosstalk when choosing expansion modalities, as even healthy donors exist on a spectrum.

While the ‘Signal Strength Model’ is a cornerstone of αβ-T-cell biology(29, 31, 32), the specific role of stimulation density in the differentiation of human peripheral blood γδ T-cells has been undetermined. Current protocols often conflate signal intensity with duration, leaving a critical gap in our understanding of how differentiation commitment and cytotoxic potential results from cumulative time or thresholds of ligand density. Our findings establish a scalable, tunable platform with control over biochemical cues for engineering γδ T-cells with improved therapeutic potential and underscore the importance of considering starting populations in adoptive cell immunotherapy design. These observations provide guidance for the rational design and optimization of stimulatory materials which could be adapted for other ligands, to accommodate individual differences among donor cells, and enable patient-specific stimulation strategies to achieve optimal therapeutic outcomes.

## Materials and Methods

### Material synthesis

Alginate microgels were fabricated via microfluidic emulsion using our previously reported procedure(19, 36). Briefly, Tz and Nb modified alginate were dissolved in DI water separately at 1–3 wt%. These solutions were mixed in microfluidic devices in situ as the dispersed phase with a final alginate concentration of 2 wt%. The fluorocarbon oil containing 1 % fluorosurfactant was used as the continuous phase. The dispersed phase was injected at an overall rate of 300 μL/h (150 μL/h for each alginate component), and the continuous phase was injected at 1000 μL/h. The emulsion was collected and left at room temperature for 24 h for complete gelation before processing with 33 % 1H,1H,2H,2H-perfluoro-1-octanol in fluorocarbon oil to break the emulsion. The collected microgels were stored at 4 °C in HEPES buffer until further use.

To modulate the stiffness of microgels, Tz and Nb functionalized alginates with different degree of modifications were used to fabricate microgels. The degree of modification of alginate was varied from 2% - 12%. The stiffness of resulting microgels were defined by characterizing the mechanical properties of bulk alginate hydrogels with the same compositions. The rheological properties of alginate hydrogels were measured using an oscillatory shear rheometer (DHR-3, TA Instruments) with an 8 mm parallel plate geometry. A time sweep test was performed at 1 Hz, 1 % strain and at 25 °C to determine the storage and loss moduli of materials.

Microgels were coated with poly(D-lysine) and azide-functionalized alginate using our published protocol(19). Briefly, microgels were resuspended in a solution of poly(D-lysine) (50–150 kDa, 0.1 mg/mL in HEPES buffer) and centrifuged at 300 rcf for 3 min. After three washes using HEPES buffer, microgels were then resuspended in a solution of azide-functionalized alginate and centrifuged at 300 rcf for 3 min. The collected microgels were washed and resuspended in HEPES buffer and used immediately after preparation. Azide-functionalized microgels were then mixed with DBCO-modified antibodies at 4 °C overnight. Microgels were washed twice times using HEPES buffer, soaked in T cell media (without cytokines) at 4 °C overnight, and washed three times with T cell media (including cytokines) to remove physically absorbed antibodies. Microgels conjugated with antibodies were used immediately after preparation for T cell activation.

### Human γδ T-cell isolation, culture, and activation

#### Human γδ T-cell isolation

Human Peripheral blood mononuclear cells (PBMCs) were obtained from Brigham and Women’s Hospital as-de-identified apheresis collars which were processed by density-gradient centrifugation using Ficoll-Paque PLUS. A fraction of fresh PBMCs was characterized by flow cytometry (see Flow Cytometry) and the remainder cryopreserved in 90% FBS / 10% DMSO. PBMCs were enriched for γδ T cells by negative selection using the Human γδ T Cell Isolation Kit (Miltenyi Biotec, 130-092-892) following the manufacturer’s instructions.

#### Human γδ T-cell culture and activation

Enriched γδ T-cells were resuspended in RPMI-1640 (Lonza, BE12-702F), supplemented with 10% heat-inactivated fetal bovine serum (Gibco,10-082-147), 1% penicillin–streptomycin, 20 mM HEPES (Gibco, 15630080), and 1x non-essential amino acids (Fisher, 11140-050), 1x sodium pyruvate (Gibco, 11360070), 55μM beta-mercaptoethanol. Cytokines were added at the following final concentrations unless otherwise indicated: 10 ng mL^-^ recombinant human IL-18 (R&D systems, 9124-IL-050); 14 ng mL^-^recombinant human IL-15 (Fisher Scientific, 10773-078). For all T-cell activations, 25k γδ T**-**cells were seeded per well. TransAct (Miltenyi Biotec, 130-111-160) was added at a 1:100 (v/v) ratio as per manufacture’s instruction. For soluble antibody-based activation, 1μg ml^-^ CD3, OKT3 clone (Fisher Scientific, 16-0037-85), and 500 ng ml^-^ CD2, RPA-2.10 clone (Fisher Scientific, 16-0029-85) were added at the start of culture and replaced simultaneously with the media as per protocol from(13). All culture conditions, including microgels, were maintained at 37 °C, 5% CO_2_, with media refreshed every 2–3 days.

### Cancer cell culture and apoptosis studies

Luciferized Human colorectal carcinoma cells (HCT116) received as a gift from the Ingber lab, were cultured with RPMI 1640(Lonza, BE12-702F) with 10% heat inactivated fetal bovine serum (Gibco,10-082-147) and 1% penicillin/ streptomycin. K562 cells were received as a gift from Lesley Kean at the Dana Farber Cancer Institute and were cultured using the same media formulation. HCT 116 cells used for the apoptosis studies were passaged at least 6 times. 4×10^3^ cells were seeded in 96-well tissue culture plastic plates overnight to allow for adhesion. K562 cells were passaged at least 6 times and 4×10^3^ cells were plated the day of assay, in suspension plates. γδ T-cells were co-cultured at a 3:1 ratio for 4 hours, then allowed to reach room temperature before addition of Promega caspase 3/7 assay reagent(Promega, G8091) as described by the manufacturer. The mixture was allowed to react for 1 hour then a BioTek Synergy H1 plate reader was used to acquire the fluorescent readout, with the peak signal used for analysis. The fold change was calculated compared to a matched non-treated cancer cell control.

### Flow cytometry

γδ T-cells were kept at 4 °C throughout immunostaining. Cells were stained with near IR live dead cell stain (Fisher Scientific, L10119) according to the manufacturer’s protocol. Cells were then blocked with FcX Fc receptor blocking solution (Biolegend, 422302) for 20 min and stained with surface protein antibodies for 30 min. Brilliant violet staining buffer (BD Biosciences, 566349) and flow cytometry staining buffer (eBioscience, 00-4222-26) were used during staining. For intracellular staining, after stimulation with the eBioscience cocktail (eBioscience, 00-4970-93) at 1x, for 1hr, Golgi plug (BD Biosciences, BDB555029) was added then after 4 hours, samples were transferred to u-bottom 96-well plates. Fixing and permeabilization carried out using the CytoFast fix and perm kit (Biolegend, NC1666067), Intracellular staining was performed per manufacturer’s instructions. Flow cytometry acquisition was carried out with the Sony ID7000 spectral cytometer. Gating was performed based on fluorescence-minus-one controls after compensation was adjusted as recommended by Sony. The complete set of antibodies used for flow cytometry are listed in Supplementary Table (1)

### Statistical analyses

Unless otherwise specified in figure legend, statistical analyses were performed on Prism GraphPad. Statistical tests used two-tailed one-way paired ANOVA, and post-hoc tests for multiple comparisons, or Student’s t-test or Wilcoxon test for comparison between two groups. Prism mixed-effects model analysis was used for datasets with missing values (where ordinary two-way ANOVA could not be performed). P < 0.05 was considered significant unless otherwise noted. Error bars represent the standard error of the mean, unless otherwise noted.

## Data, Materials and Software Availability

Study data are included in the SI appendix

## Acknowledgments

We thank Drs. Kwasi Adu Berchie, Yoav Binenbaum, and Wei-Hung Jung for their scientific inputs. We also acknowledge funding from the National Institute of Health/ National Cancer Institute grant no. 5R01CA276459-02 (FO, DM), NSF MRSEC (DMR-2011754) (JL, DM), National Science Foundation Graduate Research Fellowship (FO), Ford Foundation Fellowship (FO), MIT University Center for Exemplary Mentoring Fellowship (FO).

