## Supplementary figure (S) for "A Microgel-Based Platform for Tunable Expansion and Function of γδ T-cells"

**Affiliations**

**This PDF file includes:**

Figs. S1 to S4
Tables S1

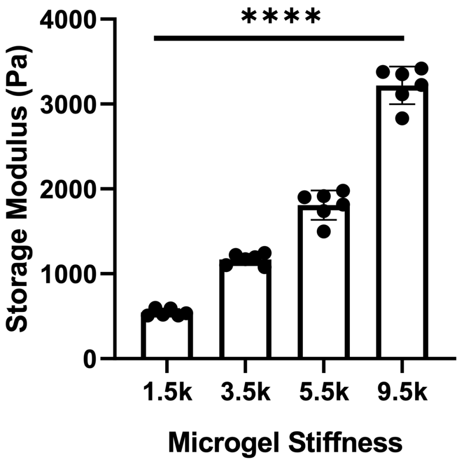


**Fig. S1. Rheological characterization of microgels with different stiffness**

**Elastic modulus of microgels can be modulated by varying the degree of substitution of norbornene and tetrazine groups on alginate polymers,** n=5 technical replicates. Statistics were performed using ordinary one-way ANOVA with Tukey’s multiple comparisons test. *P < 0.05, **P < 0.01, ***P < 0.001, ****P < 0.0001; ns, not significant.

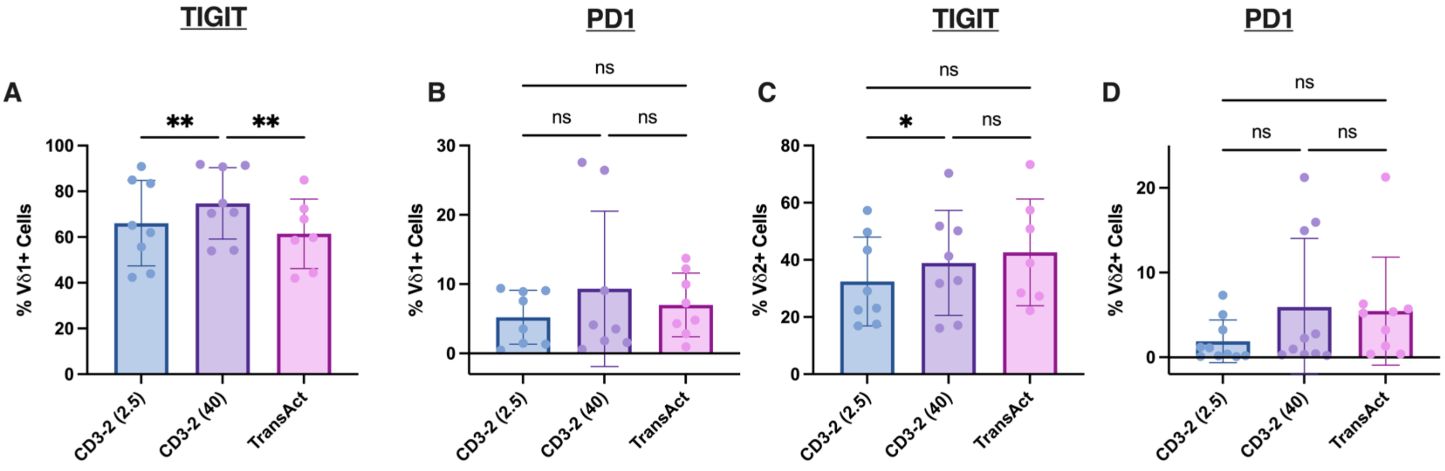


**Fig. S2. TIGIT and PD1 percentages are modulated by stimulation density**

**(A-D)** TIGT and PD1 Geometric mean fluorescence intensity for Vδ1 and Vδ2 cells respectively, across activation conditions. n=8 donors. Statistics were performed using ordinary one-way ANOVA with Tukey's multiple comparisons test. *P < 0.05, **P < 0.01, ***P < 0.001, ****P < 0.0001; ns, not significant.

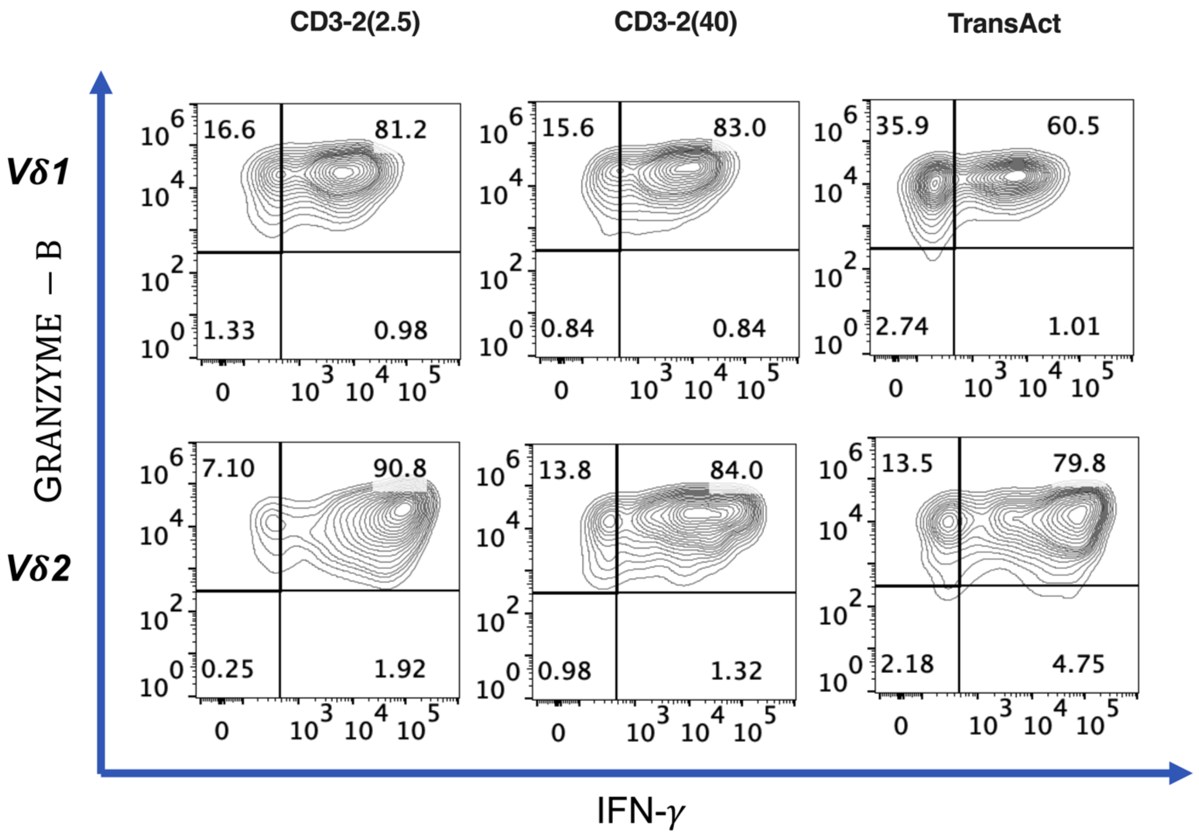


**Fig. S3. Effects of ligand density on cytotoxic potential**
Representative flow cytometry contour plots of Granzyme-B versus IFN-γ in expanded Vδ1 (top row) and Vδ2 (bottom row) γδ T-cells following culture on αCD3 functionalized microgels with αCD2 co-stimulatory molecule.

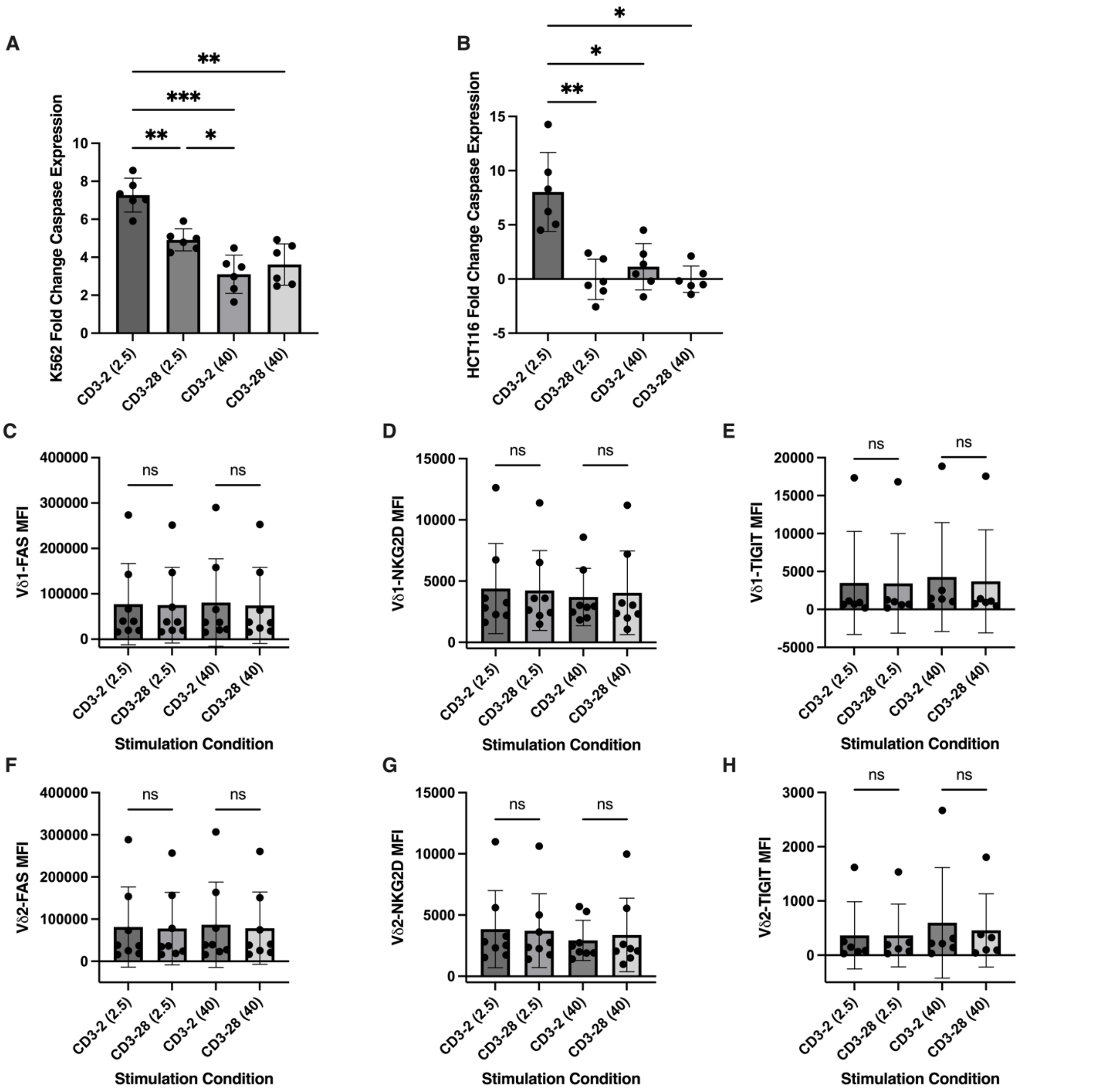


**Fig. S4. Effects of CD28 co-stimulation on activation and cytotoxicity**
(**A-B**) Fold change in caspase-3/7 signal in K562 (**A**) and HCT116 (**B**) target cells following 4 h co-culture with γδ T-cells expanded under the indicated conditions. (**C–E**) FAS, NKG2D, TIGIT geometric mean fluorescence intensity (gMFI), respectively, within the Vδ1 compartment across expansion conditions. (**F–H**) FAS, NKG2D, TIGIT geometric mean fluorescence intensity (gMFI), respectively, within the Vδ2 compartment across expansion conditions. For panels A–B n = 6 technical replicates; for panels C-D, F–G, n = 8 donors. E,H, n= 6 donors. Bar graphs show mean ± s.d. Statistics were performed using ordinary one-way ANOVA with Tukey's multiple comparisons test. *P < 0.05, **P < 0.01, ***P < 0.001, ****P < 0.0001; ns, not significant.

| Staining Antibody | Manufacturer | Cat# |
|---|---|---|
| CD3 (RB705) | BD Biosciences | 570237 |
| TCR Vδ1 (PE) | Thermo Fisher | 12-5679-42 |
| TCR Vδ2 (APC) | BioLegend | 331418 |
| CD27 (BUV395) | Thermo Fisher | 363027942 |
| CD45RA (BV510) | BioLegend | 304142 |
| PD-1 (BV605) | BioLegend | 367426 |
| NKG2D (PE-CY7) | Thermo Fisher | 25-5878-42 |
| DNAM-1 (BV785) | BioLegend | 338322 |
| NKp30 (BV711) | BioLegend | 325218 |
| CD95/FAS (PE-CY5) | Thermo Fisher | 15095942 |
| Live/Dead NIR viability dye | Thermo Fisher | L10119 |
| TIGIT (BV421) | BioLegend | 372710 |
| TCRγδ (PE/Dazzle) | BD Biosciences | 331226 |
| IFNγ (FITC) | BD Biosciences | 554551 |
| Granzyme B (PE-Texas Red) | Invitrogen | GRB17 |
| TNFα (BV421) | Thermo Fisher | 404-7349-42 |

**Table S1. Antibodies used for experiments**
List of antibodies used for flow cytometry, with manufacture’s catalog number provided. The antibodies were used at 1:60 dilution, except for Live Dead NIR which was 1:1000 dilution.
